# Allele-specific epigenetic activity in prostate cancer and normal prostate tissue implicates prostate cancer risk mechanisms

**DOI:** 10.1101/2021.06.02.446619

**Authors:** Anamay Shetty, Ji-Heui Seo, Connor A. Bell, Edward P. O’Connor, Mark Pomerantz, Matthew L. Freedman, Alexander Gusev

## Abstract

**Background:** Genome-wide association studies of prostate cancer have identified >250 significant risk loci, but the causal variants and mechanisms for these loci remain largely unknown. Here, we sought to identify and characterize risk harboring regulatory elements by integrating epigenomes from primary prostate tumor and normal tissues of 27 patients across the H3K27ac, H3K4me3, and H3K4me2 histone marks and FOXA1 and HOXB13 transcription factors.

**Results:** We identified 7,371 peaks with significant allele-specificity (asQTL peaks). Showcasing their relevance to prostate cancer risk, H3K27ac T-asQTL peaks were the single annotation most enriched for prostate cancer GWAS heritability (40x), significantly higher than corresponding non-asQTL H3K27ac peaks (14x) or coding regions (14x). Surprisingly, fine-mapped GWAS risk variants were most significantly enriched for asQTL peaks observed in tumors, including asQTL peaks that were differentially imbalanced with respect to tumor-normal states. These data pinpointed putative causal regulatory elements at 20 GWAS loci, of which 11 were detected only in the tumor samples. More broadly, tumor-specific asQTLs were enriched for expression QTLs in benign tissues as well as accessible regions found in stem cells, supporting a hypothesis where some germline variants become reactivated during/after transformation and can be captured by epigenomic profiling of the tumor.

**Conclusion:** Our study demonstrates the power of allele-specificity in chromatin signals to uncover GWAS mechanisms, highlights the relevance of tumor-specific regulation in the context of cancer risk, and prioritizes multiple loci for experimental follow-up.

## Background

Genome-wide association studies (GWAS) have identified >250 known risk loci for prostate cancer (PrCa) (Schumacher et al. 2018; Conti et al. 2021; Dadaev et al. 2018). However, understanding risk mechanisms remains challenging because most associations map to noncoding regions and likely influence target gene transcription by modifying regulatory element binding in specific contexts. Several techniques have been developed to connect common noncoding variants to target genes through integration or colocalization with expression quantitative trait loci (eQTLs) (Gusev, Ko, et al. 2016; Gamazon et al. 2015; Barbeira et al. 2018). For example, a recent Transcriptome-wide Association Study (TWAS) of prostate cancer used eQTL signals to identify 217 putative target genes at 84 independent loci (Mancuso et al. 2018). However, such methods do not seek to identify the regulatory element driving the GWAS association and typically cannot identify a putative mechanism for the majority of loci. Specific to the cancer context, it remains unknown whether risk mechanisms are better captured by QTLs in normal tissues, which may miss the appropriate cell type or context of the mechanism, versus QTLs in tumors, which may better reflect the cell of origin (Polak et al. 2015).

As with eQTL colocalization, studies of QTLs associated with chromatin activity (cQTLs) present an opportunity to understand the link between GWAS SNPs via their effect on *cis*-regulation elements. Recent studies in healthy individuals have identified cQTLs for a variety of epigenetic features such as DNA accessibility (Gate et al. 2018), transcription factor (TF) binding (Maurano et al. 2015), and histone marks (Waszak et al. 2015; Grubert et al. 2015). The *cis*-regulatory landscape has been shown to be rich with co-regulation and co-operative binding, making it harder to connect non-coding disease variants to mechanism (Link et al. 2018). A key limitation to cQTL studies has been the large sample size needed to detect such variants. This is because *cis*-genetic effects tend to be small compared to *trans*-regulatory effects (X. Liu, Li, and Pritchard 2019) and technical variation between individuals.

Recent advances in molecular sequencing technology have enabled the identification of QTLs in a single individual through measurements of allelic imbalance. Allelic imbalance quantifies the difference in reads mapping to a heterozygous variant within an individual, and is indicative of a cis-regulatory effect on one of the haplotypes. Methods based on allele-specific reads benefit from the ability to remove *trans*--effects by comparing alleles in the same environment, increasing power to detect *cis*-effects from *in vivo* samples. Individual-level signals can then be aggregated across individuals to identify common germline regulatory variants. This technique has been applied to detecting QTLs (Kumasaka, Knights, and Gaffney 2016; van de Geijn et al. 2015) and gene-by-environment interactions (Knowles et al. 2017) in healthy samples. Recently, we extended this approach to identify differences in allelic imbalance across many individuals between their tumor/normal cell states (Gusev et al. 2019). Such state specific QTLs may highlight tumor-specific *cis*-regulatory mechanisms or interactions that would be difficult to observe in normal tissues or healthy controls

In this work, we used germline allelic imbalance in epigenomes from prostate tumors and matched normal tissue to describe the mechanisms of genetic risk for prostate cancer. We identified thousands of allelically imbalanced *cis*-regulatory elements at histone marks and transcription factors (HMTFs), including hundreds of variants that with significant changes in their regulatory effect between the normal/tumor cell states. These *cis*-regulatory elements were enriched for eQTLs, GWAS associations, cell-type specific accessible regions, and complex patterns of transcription factor binding, shedding light on mechanisms of prostate cancer risk and tumorigenesis.

## Results

### Identification of allelic imbalance in prostate epigenomes

We collected ChIP-seq data from 27 patients in primary prostate cancer tumor samples and adjacent normal prostate tissue (Pomerantz et al. 2020, 2015). We assayed epigenomes from three histone marks (HM) and two transcription factors (TF) which have previously been implicated in prostate cancer development (Pomerantz et al. 2015; Whitington et al. 2016): H3K27ac (n=26), H3K4me3 (n=3), H3K4me2 (n=3); *FOXA1* (n=5), *HOXB13* (n=5) (Supplementary Table 1). Peaks were called using the MACS2 software, and mapping bias was corrected using the WASP pipeline (van de Geijn et al. 2015) (see Material and Methods). A principal components analysis revealed no clear differences in ChIP-seq signal between tumor and normal peaks (Supplementary Figure 1) in contrast to previous studies in other cancers (Gusev et al. 2019).

We tested each peak for allelic imbalance across all individuals using a haplotype-based test implemented in the stratAS algorithm (Gusev et al. 2019). Unlike some conventional allelic imbalance models, this is a test for consistent allelic effect across multiple individuals, and thus captures QTLs in the sampled population. For each individual, reads were first summed across all phased heterozygous variants within a ChIP-seq peak to quantify haplotype-specific read counts. For each peak, all individuals with haplotype-specific read counts were then tested for consistent allelic imbalance at each SNP within 100kb of the peak center using a beta-binomial test (Figure 1). The test was run three times using different samples: across all samples regardless of cell state (“T+N”); across all normal tissue samples (“N”); across all tumor samples (“T”). In addition, a beta-binomial likelihood ratio test was used to identify those SNPs with a significant difference in allelic fraction between normal and prostate cancer tissue (“TvsN”). These categories are termed as ‘states’ and, in combination with the five histone marks and transcription factors (“HMTFs”), yielded 20 independent HMTF-state categories of ChIP-seq peak. After testing, a 10% FDR threshold was applied to each HMTF-state group to identify significantly imbalanced SNPs. We refer to those peaks which were associated with significant SNPs as “asQTL peaks”. Those peaks which were tested (i.e. contained heterozygous variants and satisfied the minimum coverage requirements to be testable) but not significantly associated with asQTLs were termed ‘balanced’ peaks.

**Figure 1:**
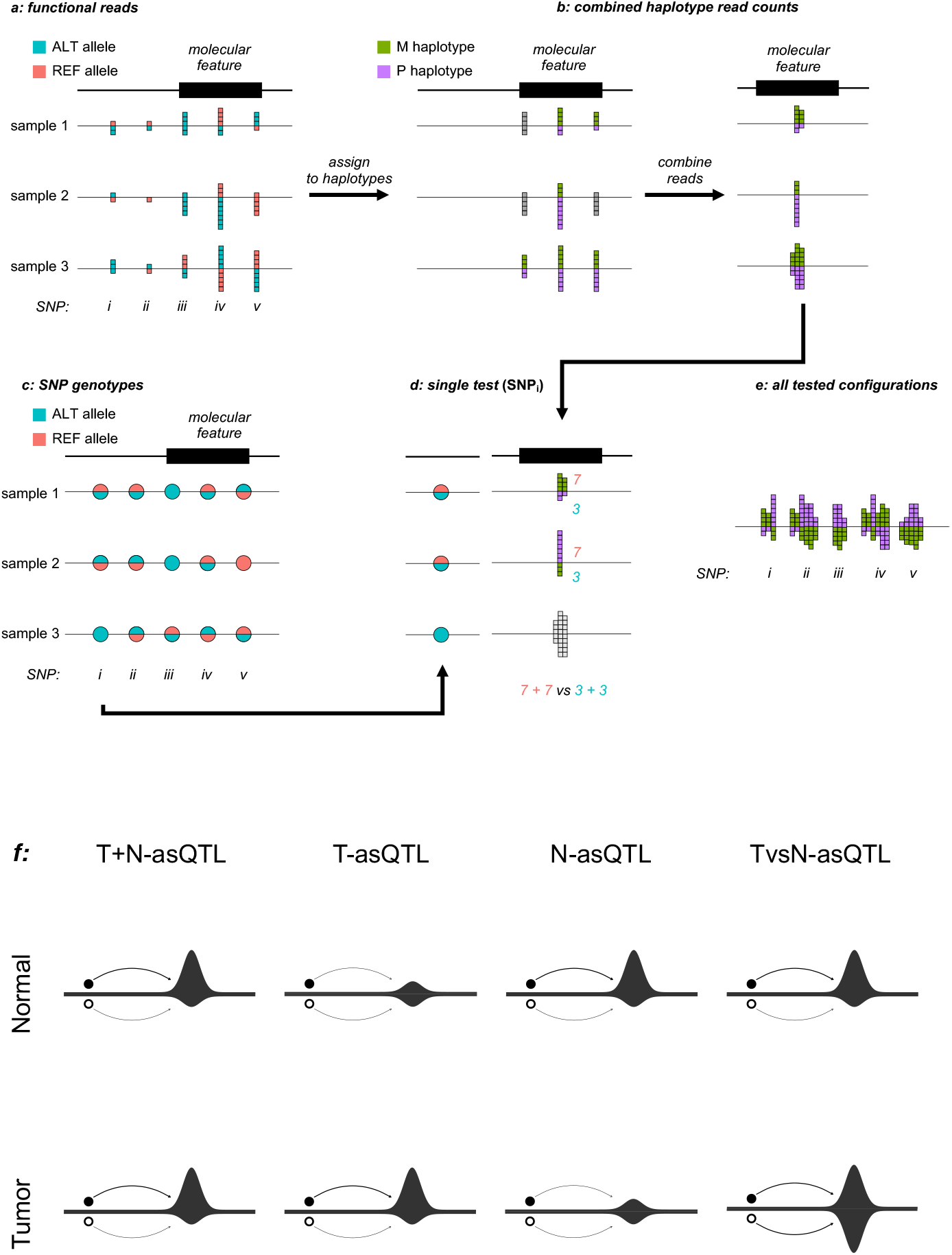
The stratAS framework: a method to leverage allelic imbalance to identify causal variants stratAS takes in two inputs - a matrix of allelic read counts by SNP site and sample (a) and the genotypes by SNP site and sample (c). The algorithm iterates over each molecular feature by collating the reads of all heterozygous SNPs within a molecular feature to develop the sum of reads on either haplotype for each sample (b). This haplotype allelic imbalance is converted to a genotype allelic imbalance by the genotype data from step (c), where the allelic read counts for the feature is summed across samples, for each SNP (i-v shown) in the testing window. The allelic read count distributions are then assessed via a beta-binomal test (e) to determine if the allelic fraction is sufficiently polarised between a variant allele on one haplotype and a reference allele on another haplotype. Figure (f) provides a graphic showing how allelic imbalance arises in each of the four states of peaks. In T+N, allelic imbalance – shown as the difference in read count between the haplotype associated with the variant allele (black circle) compared to the reference allele (white circle) – is present in both tumor and normal samples. In T-asQTL and N-asQTL, allelic imbalance is only present in the respective sample types. In TvsN-asQTL peaks, there is a significant change in the allelic imbalance between the tumor and normal states – either because one state is balanced but contains a high read count (as tumor is depicted as being in the example) or the imbalance in is in the opposite direction (not depicted).

### Primary prostate tissues harbor thousands of asQTL peaks

We identified 7,371 unique asQTL peaks (at 10% FDR) across the five HMTFs tested (Table 1) out of 525,634 peaks tested by stratAS: 5,975 T+N peaks; 2,221 N peaks; 4,006 T peaks; and 1,606 TvsN peaks. There was a significant correlation between the number of allelic reads in a sample and the number of asQTL peaks detected within the sample (Spearman’s rank correlation=0.523, p<0.01) as this is the primary determinant of statistical power. Although *FOXA1* and *HOXB13* TF activity was measured in fewer samples, these TFs had a greater yield of asQTL peaks per reads per base pair than the histone marks (p<0.01 in a linear model) (Figure 2A).

**Figure 2:**
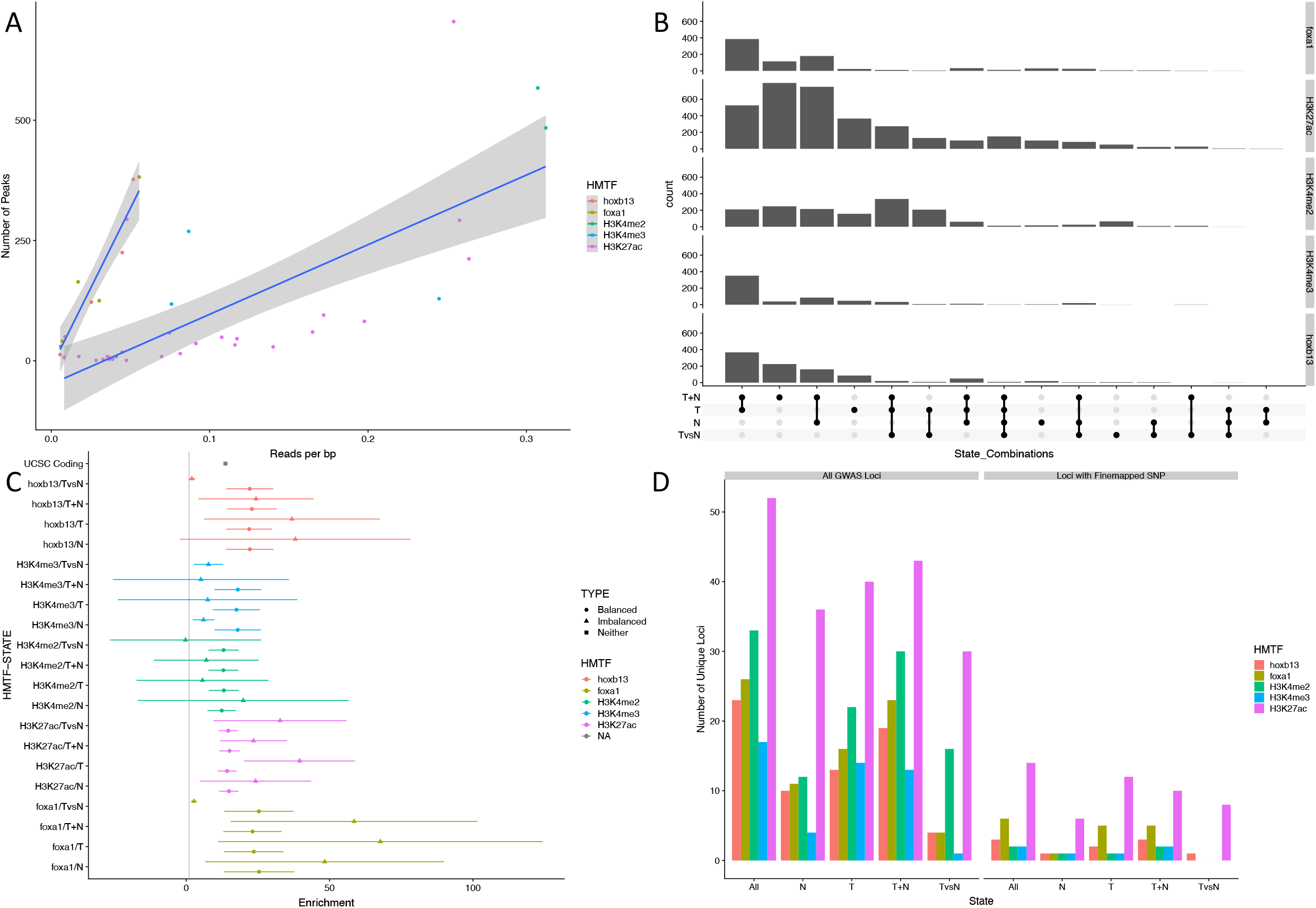
Descriptive summary of asQTL peaks and enrichment for key features A, a scatterplot of the read density per sample and the number of peaks per sample. Regression lines show the estimated slopes of peaks against enrichment for the transcription factors (*HOXB13/FOXA1*) against histone marks (H3K27ac, H3K4me3, H3K4me2). B, an upset plot between the 4 different state categories to show the number of overlapping peaks. C, the estimates from LD-score regression. Absent results are those which failed to converge. D, the number of GWAS loci within certain HMTF-state categories and loci within these categories which contain a fine-mapped SNP.

There was significant overlap in asQTL peaks across states within a given HMTF as well as across different HMTFs. Within a given HMTF, significant asQTL peaks were frequently observed in multiple states (Figure 2B). Taking H3K27ac peaks as an example, a minority of asQTL peaks were only significant in a single state: 366 (24%) T-asQTL peaks, 100 N-asQTL peaks (8%) and 51 TvsN-asQTL peaks (7%). For HOXB13 and FOXA1, six (11%) and seven (10%) TvsN-asQTL peaks were unique to TvsN-asQTL, respectively. TvsN peaks were evenly divided between those with imbalance in favour of the variant in normal tissue samples and those in prostate cancer samples (Supplementary Figure 2).

Across HMTFs, asQTL SNPs often exhibited significant correlation in allelic fraction (the measure of regulatory effect size) when measured in different HMTFs (Supplementary Table 2). The largest positive correlations were observed between *FOXA1* and *HOXB13* (ρ=0.75, p<0.01), H3K27ac and *HOXB13* (ρ=0.66, p<0.01); whereas both *FOXA1* and *HOXB13* exhibited negative correlations with H3K4me3 (ρ=-0.14, p<0.01; ρ=-0.24, p<0.01 respectively). Similarly, asQTL peaks were generally closer than random to other ChIP-seq peaks, suggestive of coordinated histone activity across nearby marks under cis regulatory control (Kumasaka, Knights, and Gaffney 2019) (Supplementary Table 3).

### asQTL peaks localize PrCa GWAS heritability

PrCa risk loci are enriched in epigenomic features (Gusev, Shi, et al. 2016) and we hypothesized this enrichment would be stronger in asQTL peaks that are under cis regulation. We evaluated risk variant enrichment using data from a recent large-scale PrCa GWAS study in >140,000 samples (Schumacher et al. 2018) and multiple enrichment techniques.

First, we partitioned PrCa GWAS heritability across regulatory peaks from our data using stratified LD-score regression (S-LDSC) (Supplementary Table 4, Figure 2C; see Materials & Methods). S-LDSC identifies enrichments in polygenic GWAS effect sizes within a given annotation, while accounting for co-incidental background enrichment from a “baseline” model of broad functional features. Surprisingly H3K27ac T-asQTL peaks exhibited the most significant enrichment (40x s.e. 9.9) which was significantly higher than that of balanced H3K27ac peaks (P=0.01 for difference by z-test). T-asQTL, T+N-asQTL and N-asQTL *FOXA1* peaks had the largest enrichments by magnitude (68x, 59x, 48x respectively) but were not significantly different from their respective balanced peak categories, likely due to insufficient power for these small annotations (Reshef et al. 2018). For comparison, coding regions were 14x (s.e. 4.1x) enriched and evolutionarily conserved regions were 18x (s.e. 4.1x) enriched. Within balanced peaks, enrichments were significant across all HMTFs but generally weaker, with the most significant enrichment from T+N H3K27ac peaks (15x s.e. 1.8) and highest magnitude of enrichment from N *FOXA1* peaks (25x s.e. 6.3). A SNP in an imbalanced regulatory region is thus expected to explain more PrCa heritability than even a coding variant.

Second, we measured the enrichment of fine-mapped PrCa GWAS SNPs which have a high probability of being causal (Dadaev et al. 2018). For each of the 20 HMTF-state combinations of peaks, we calculated the ratio of GWAS SNPs per base pair in all asQTL peaks, to GWAS SNPs per base pair in the same number of randomly sampled balanced peaks. We term this the enrichment ratio, 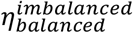. This comparison to randomly sampled balanced peaks was a stringent baseline that specifically estimates the enrichment beyond the general enrichment of functional variants in epigenetically active regions. We assessed significance by performing multiple samplings of balanced peaks and estimating an empirical p-value relative to this null distribution (see Materials & Methods).

Of the 20 HMTF-state combinations, 3 were significantly enriched for GWAS fine-mapped SNPs compared to balanced peaks (empirical p-value < 0.05): H3K27ac T-asQTL peaks (8.5x, empirical p-value<0.001), H3K27ac TvsN-asQTL peaks (8.3x, p<0.001) and H3K27ac T+N-asQTL, peaks (5.5x, p<0.001) (Figure 3, Table 2). All three significant enrichments involved measurements from tumors, and H3K27ac N-asQTL peaks were not significantly enriched (1.83x, p=0.20). As a check, we carried out the same enrichment procedure with a background of randomly selected genomic intervals instead of balanced peaks (termed 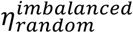), which yielded substantially higher enrichments across all categories as expected, including 38.2x for TvsN-asQTL H3K27ac peaks and 10.0x for N-asQTL H3K27ac peaks (Supplementary Figure 3, Supplementary Table 5). H3K27ac asQTLs in tumors and specific to tumors are thus significantly more likely to harbor causal PrCa variants than random H3K27ac peaks.

**Figure 3:**
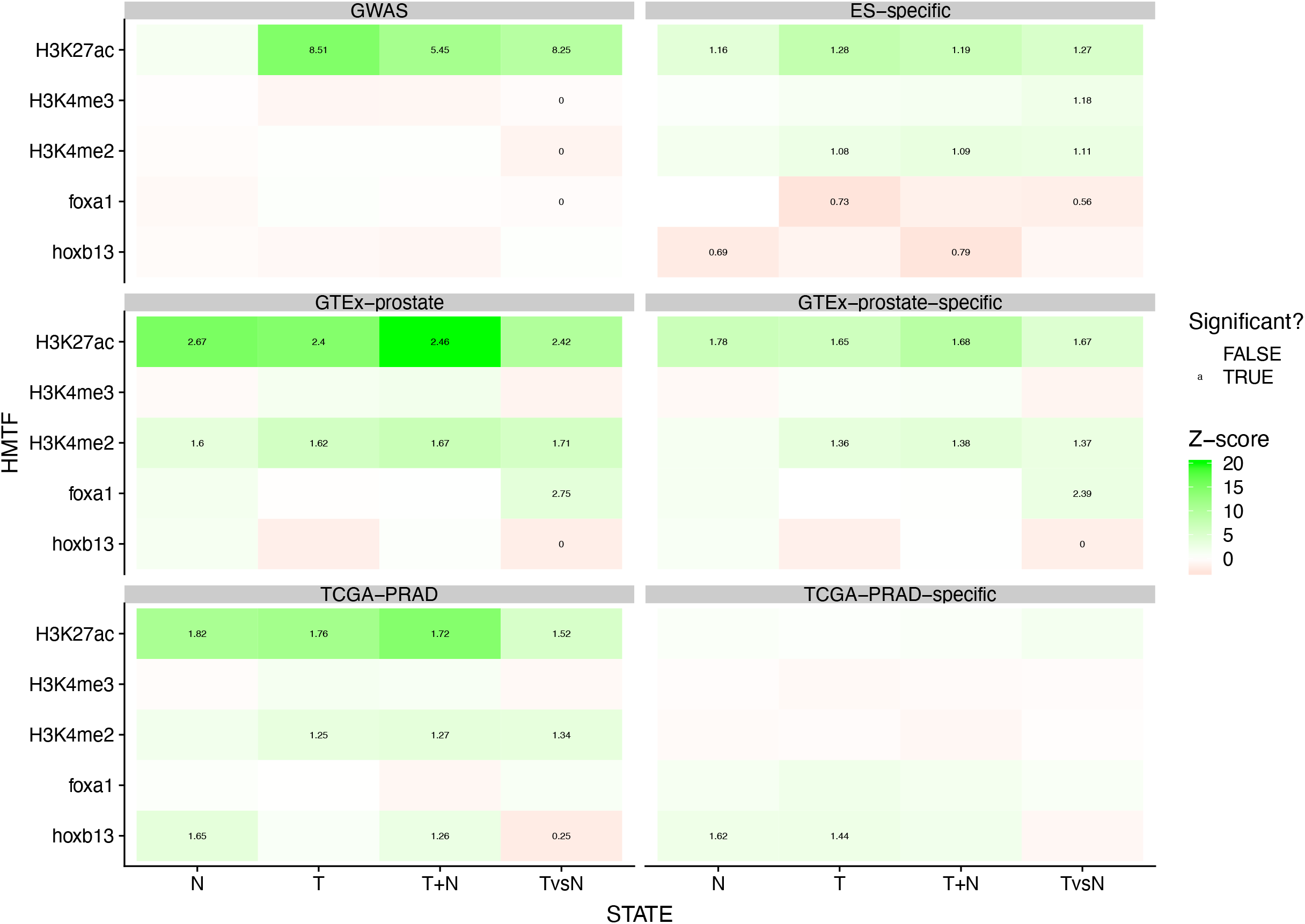
Enrichment of asQTL peaks for key genomic and epigenomic features The enrichments of the 20 different HMTF-state categories for various genetic and epigenetic markers. Colour denotes the z-score associated with the enrichment: green indicating a positive z-score and red indicating a negative z-score. Boxes labelled with a number - indicating the fold enrichment - denote that the category was enriched or depleted for the feature of interest with an empirical p-value less than 0.05.

### asQTL peaks implicate specific regulatory mechanisms at PrCa GWAS loci

We quantified how many GWAS loci contained a fine-mapped SNP within an asQTL peak, and thus had a putative regulatory mechanism. We defined 71 contiguous regions within 1MB of a genome-wide significant variant. In total, 20 out of 71 genome-wide significant regions contained fine-mapped SNPs that overlapped an asQTL peak (Figure 2D). For comparison, a recent TWAS of the same GWAS data using gene expression from 483 prostate tumor samples (20x larger than our study) identified a significant association at 12/71 GWAS loci (Mancuso et al. 2018). Across states, 11/20 loci overlapped T-asQTL or TvsN-asQTL peaks but did not overlap N-asQTL peaks, highlighting the utility of profiling tumors and suggesting a sizeable fraction of risk mechanisms may be unobservable in normal tissues of comparable sample size, consistent with our previous tumor-specific heritability enrichments and previous hypotheses (Mancuso et al. 2018; Gusev, Shi, et al. 2016; Geeleher et al. 2018). Across HMTFs, H3K27ac asQTL peaks overlapped fine-mapped SNPs in the greatest number of GWAS loci (14), followed by FOXA1 (6) and HOXB13 (3). TF asQTL peaks overlapped 8 unique loci, of which 4 did not overlap HM asQTL peaks (Supplementary Table 6). These findings highlight the utility of TF ChIP-seq to identify potential risk mechanisms not implicated by broader histone marks (Benaglio et al. 2019).

As a specific example, we investigated the *TMPRSS2* risk locus, which acquires a somatic *TMPRSS2-ERG* fusion in >50% of prostate tumors in men of European ancestry (Tomlins et al. 2005), and is suspected to harbor germline-somatic interactions (Emami et al. 2019). The locus contained a distal *HOXB13* T+N-asQTL peak overlapping three fine-mapped variants in close proximity, which was flanked by H3K27ac T-asQTL peaks (Figure 4). These three variants resided within an active tumor-specific Androgen Receptor (AR) binding site (Pomerantz et al. 2020), further supporting their role as the likely causal variants. The locus additionally contained a group of 9 TvsN-asQTL H3K27ac peaks ~10kb downstream and overlapping the *TMPRSS2* promoter, possibly co-active with the distal regulatory element. These overlaps with tumor and tumorspecific peaks further support the hypothesis of germline-somatic interactions at this locus and showcase the utility of asQTLs to localize the putative causal mechanism.

**Figure 4:**
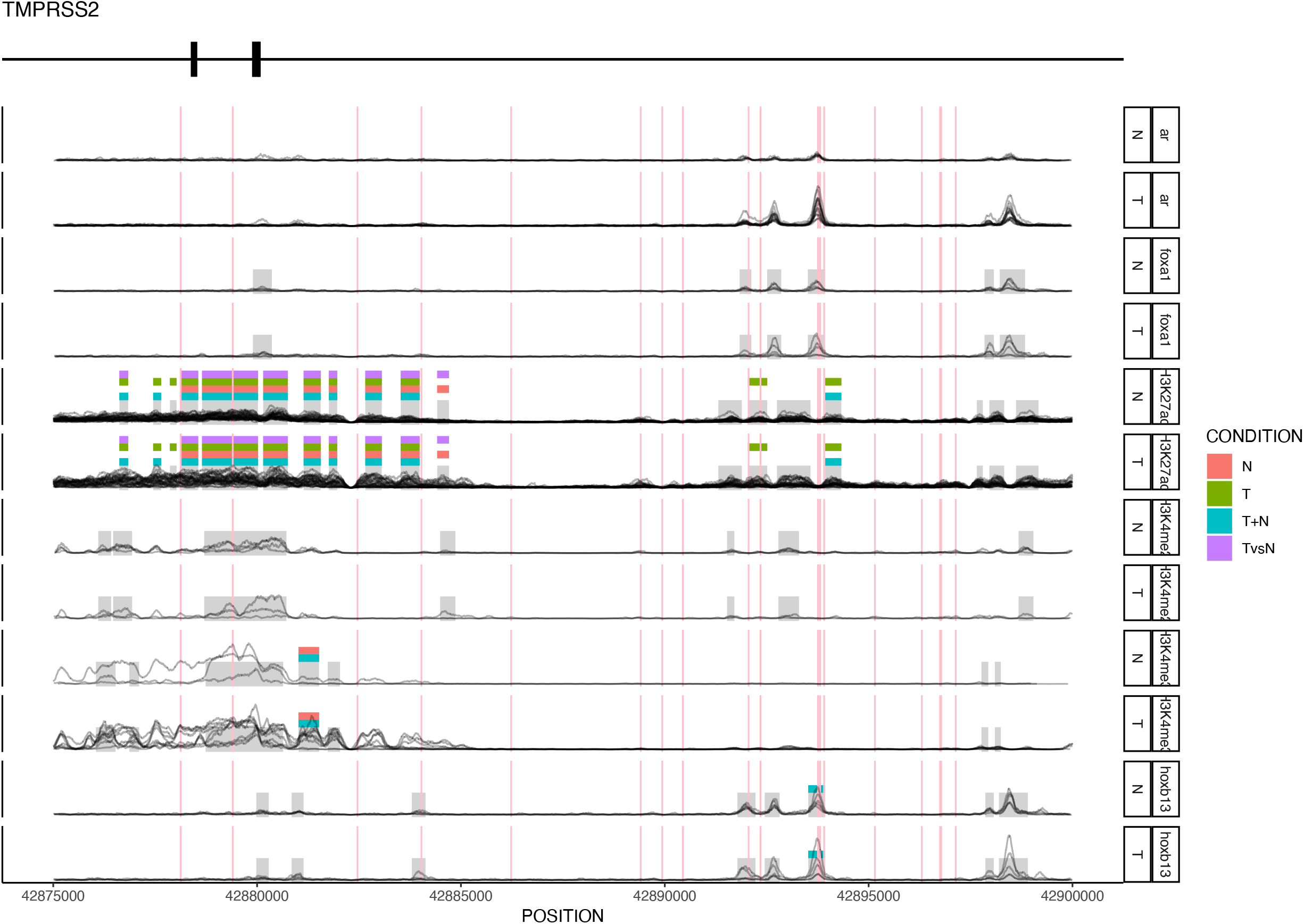
Genomic loci annotated with asQTL peaks This graphic represents the ChIP-seq reads for 6 epigenetic marks for the *TMPRSS2* locus. Grey rectangles indicate ChIP-seq peaks indicated by the *MACS2* algorithm, whilst coloured rectangles above indicate whether these peaks are imbalanced, and in what cell states. Pink vertical lines indicate the presence of GWAS fine-mapped SNPs.

### asQTL peaks contain lineage and tumor specific eQTLs

We next sought to characterize the transcriptional activity at asQTL peaks by integrating them with independent expression QTL (eQTL) data. We reasoned that asQTL peaks were more likely to harbor variants with causal effects on transcription of nearby genes and should thus be enriched for eQTLs in relevant tissues. First, we calculated the enrichment of HMTF-state combinations for eQTLs with the same procedure used to calculate enrichment of PrCa GWAS SNPs. We used eQTLs from the GTEx Consortium in prostate tissue as representative healthy prostate eQTLs (referred to as “GTEx-prostate eQTLs”), and eQTLs from prostate adenocarcinoma (PRAD) samples from TCGA as representative of PrCa tumor eQTLs (TCGA-PRAD) (Figure 3, Supplementary Table 7). We again conservatively estimated eQTL enrichment within asQTL peaks compared to balanced peaks 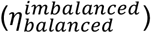.

In total there were 10 HMTF-state combinations with nominally significant 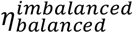 (p<0.05) for GTEx-prostate eQTLs and 10 for TCGA-PRAD eQTLs. All asQTL H3K27ac peak categories were significantly enriched for GTEx-prostate eQTLs and TCGA-PRAD eQTLs. T-asQTLs were similarly enriched in both GTEx prostate and TCGA-PRAD, showing that T-asQTLs capture broad germline effects on transcription from both healthy and cancer tissues. Across all features, TvsN-asQTL *FOXA1* peaks had the greatest enrichment for GTEx-prostate eQTLs (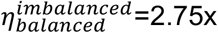, p<0.01) whilst N-asQTL H3K27ac peaks had the greatest enrichment for TCGA-PRAD eQTLs (1.82x, p<0.01). Interestingly, TvsN-asQTL *HOXB13* peaks exhibited a significant depletion in GTEx-prostate eQTLs. (p<0.01; Figure 3, Supplementary Table 7), suggestive of novel tumorspecific regulatory variants. As a sanity check, we again estimated the enrichment relative to random regions in the genome (rather than active elements; 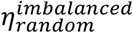), finding significant enrichments for 19/20 and 17/20 HMTF-state combinations in TCGA-PRAD and GTEx-prostate eQTL respectively (empirical p<0.05) (Supplementary Figure 3, Supplementary Table 5).

We quantified the tissue-specificity of asQTLs by comparing them to eQTLs identified in non-prostate tissues from the GTEx and TCGA studies (as there are currently no large scale, multitissue chrom-QTL studies). We computed the ratio of GTEx-prostate eQTLs enrichment to GTEx-non-prostate eQTL enrichment, and likewise for TCGA-PRAD relative to TCGA-non-PRAD. By using the enrichment relative to non-prostate tissues as a background, we generated an enrichment ratio of the form 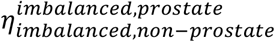 and 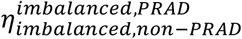 (see Materials & Methods). In GTEx, nine tissue-specific enrichments were significant at a threshold of empirical p<0.05, the greatest enrichment being TvsN-asQTL *FOXA1* peaks (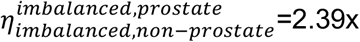, empirical p=0.028) (Figure 3, Supplementary Table 7). All H3K27ac three H3K4me3 categories exhibited a significant 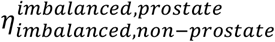. In TCGA, two were significant: N-asQTL *HOXB13* 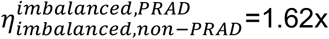, p=0.05) and T-asQTL *HOXB13* peaks (1.44x, p=0.03 respectively). These tissue-specific enrichments highlight asQTLs as broadly enriched for prostate specific effects on transcriptions, and asQTLs in *HOXB13* as being enriched for prostate tumor specific effects.

### asQTL peaks show evidence of somatic dedifferentiation

Studies of prostate tumor epigenomes have identified the phenomena of somatic dedifferentiation, whereby the tumor epigenome becomes less specific to the normal cell of origin (Brawn 1983; Pomerantz et al. 2020). We hypothesized that dedifferentiated regulatory elements harboring a germline regulatory variant could thus become activated in the tumor and detectable as T-asQTLs (if the regulatory element is newly active in the tumor) or TvsN-asQTLs (if the regulatory element is active but poised in the normal) (Figure 5). We would thus expect T-/TvsN-asQTLs to be enriched for dedifferentiated regions. To test this, we used H3K27ac peaks from ROADMAP in embryonic stem (ES) cells as a surrogate for the dedifferentiated epigenetic state (see Materials & Methods). We then calculated an enrichment of the same form as the tissuespecific scores, using H3K27ac peaks from fetal prostate cells as a baseline. As with the tissuespecific scores, this allows us to compare enrichment relative to balanced features and our background of interest. This creates an enrichment coefficient of the form 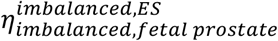.

**Figure 5:**
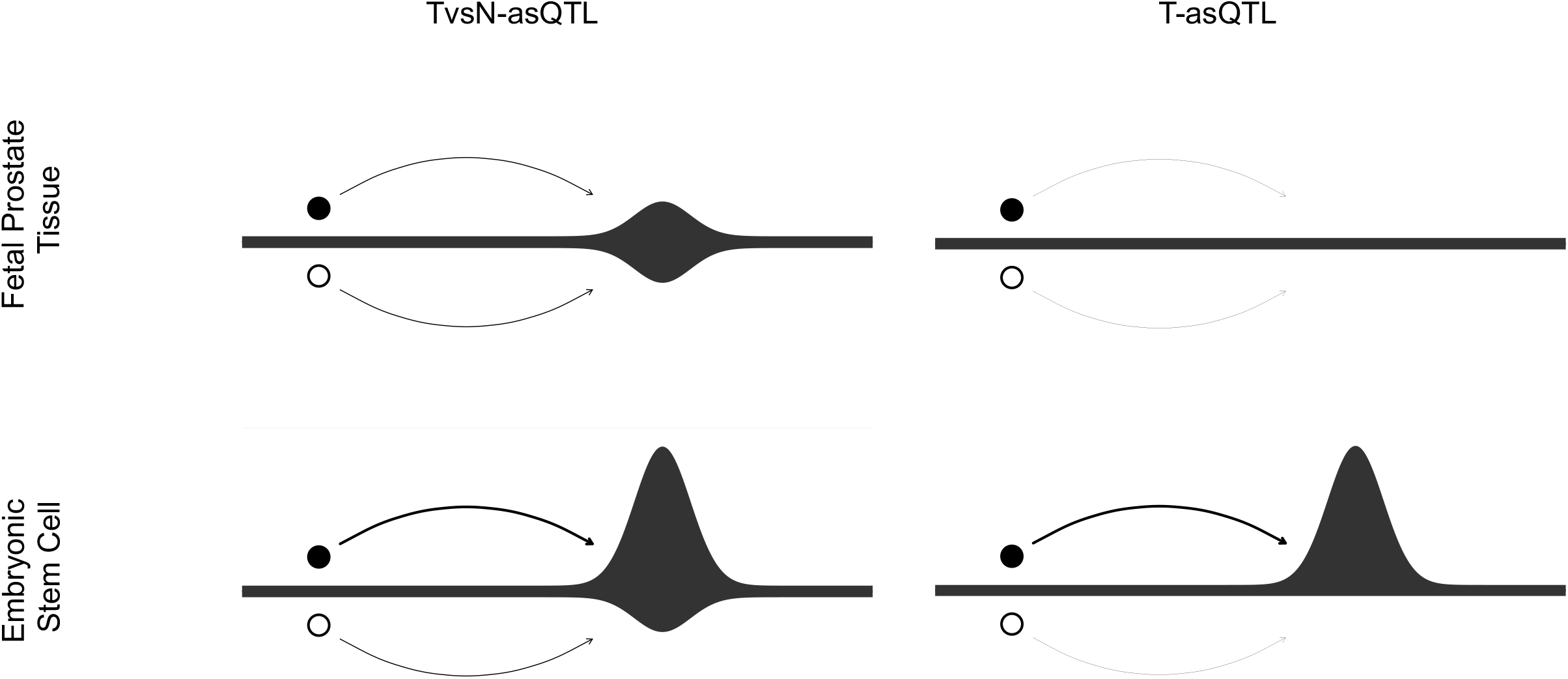
Graphical illustration of de-differentiation The graphic shows two potential circumstances where we can see allelic imbalance between the fetal prostate cells and embryonic stem cells. In T-asQTL peaks, this is due to the presence of activity in the haplotype containing the variant and no activity in the reference haplotype: in TvsN-asQTL peaks, there is a change in allelic fraction in the background of constitutive binding.

We observed significant enrichment for dedifferentiated regions across all H3K27ac asQTL peak states, with the strongest in T-asQTL (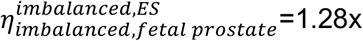, p<0.01) and TvsN-asQTL (1.27x, p<0.01) peaks (Figure 3, Supplementary Table 7). In contrast, HOXB13/FOXA1 asQTL peaks exhibited significant dedifferentiation depletion, ranging from 0.56x (p=0.0492) for TvsN-asQTL *FOXA1* peaks to 0.79x (p=0.0028) for T+N-asQTL *HOXB13* peaks (Figure 3, Supplementary Table 7), consistent with prostate lineage specificity of these two transcription factors. Only N-asQTL *FOXA1* peaks were significantly enriched (1.21x, p=0.03). Combining T-asQTL and TvsN-asQTL peaks did not change the significance of the results compared to considering each category separately, apart from T+TvsN-asQTL *FOXA1* peaks, which became non-significantly depleted (0.64x, p=0.0616) (Supplementary Figure 4). This enrichment lends support to our hypothesis that germline regulatory variants may become “reactivated” in the tumor through dedifferentiation.

### asQTL peaks implicate novel cancer-specific transcription factors and extensive cooperative binding

Motivated by the observation that asQTLs may be capturing tumor-specific regulatory mechanisms, we applied our enrichment method to TFs to nominate novel TFs that may be relevant to prostate tumorigenesis. We used publicly available ChIP-seq measurements on 25 transcription factors and epigenetic marks assayed in the LNCaP cell line and processed them using the Cistrome pipeline (T. Liu et al. 2011). Enrichment of TF activity at imbalanced asQTL peaks was widespread, with 219 out of 500 LNCaP-TF/HMTF-state combinations exhibiting significant enrichment at an FDR of 1% (Figure 6A, Supplementary Table 8). Notably, *AR* was significantly enriched in H3K27ac T-asQTL (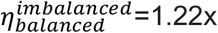, p<0.001) and TvsN-asQTL (1.22x, p<0.001), consistent with the known phenomena of AR reprogramming during transformation (Pomerantz et al. 2015).

**Figure 6:**
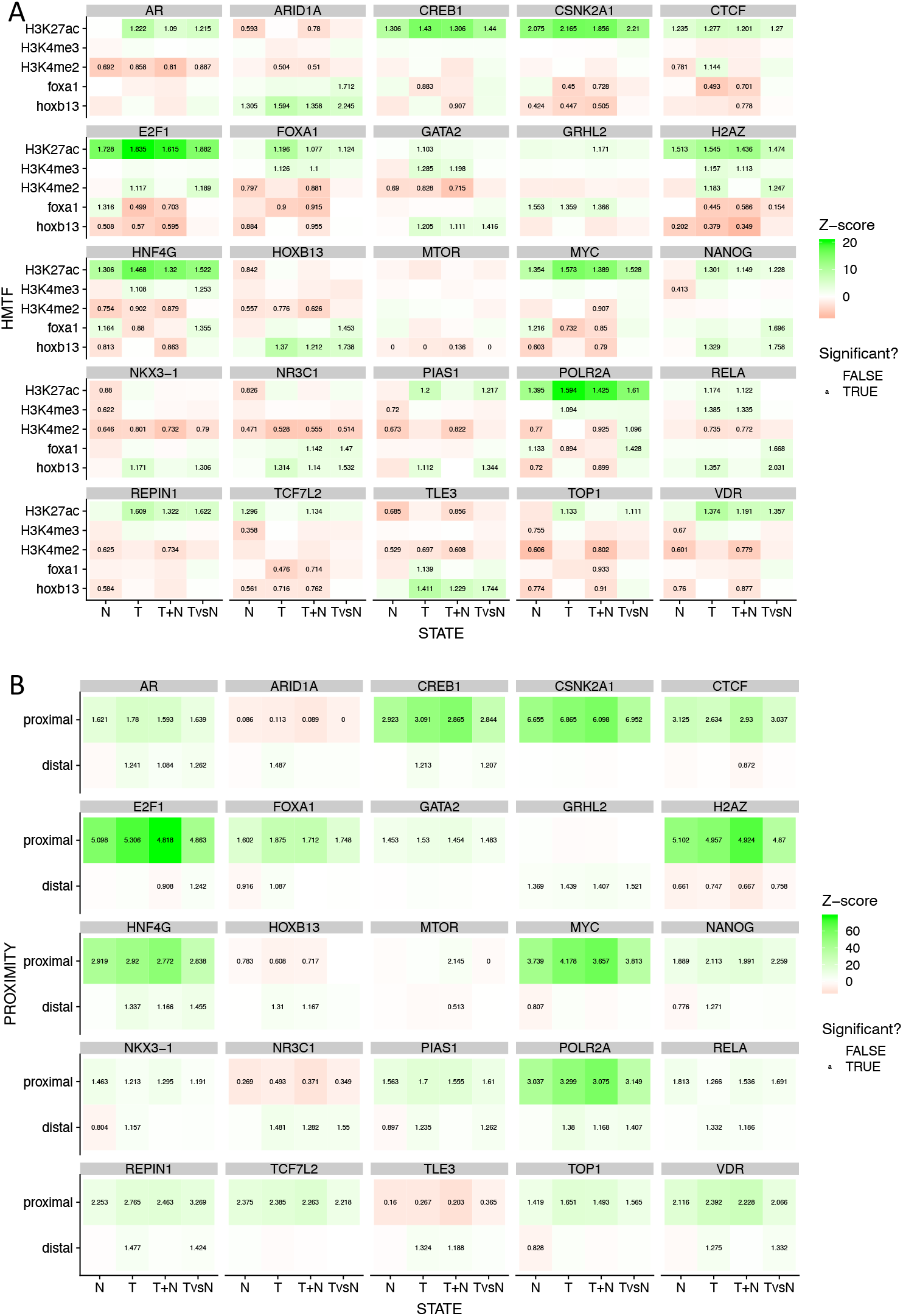
Enrichment of asQTL peaks for transcription factors and other histone features A, Enrichment of the 20 different HMTF-state categories of asQTL for 25 transcription factors and histone features from ROADMAP. B, Enrichment of h3k27ac peaks by STATE and proximity (defined by whether it lies greater or less than 1kb) to a TSS for 25 transcription factors and histone modifications. Fold enrichment in both is shown for HMTF-state combinations at an FDR of less than 1%.

Given the breadth of significant enrichment, we further stratified the H3K27ac peaks to those within 1kb of a transcription start site (“proximal”, and likely acting directly via the promoter) and those that were not (“distal”, and likely acting through enhancer/repressor elements looping into a target promoter) and re-evaluated enrichment of TFs at asQTL peaks within each sub-category (Figure 6B, Supplementary Table 9). Serving as positive controls, canonical PrCa TFs were enriched at distal T-asQTLs but not distal N-asQTLs: *AR*, *FOXA1, HOXB13*, and *NKX3-1 (Bhatia-Gaur et al. 1999)*. In addition to these canonical factors, multiple novel TFs were significantly enriched in distal T or TvsN asQTL-peaks, which we broadly classified into three groups:

1. TFs that collaborate with the above canonical TFs: *CREB1*, which co-localizes with FOXA1(Sunkel et al. 2016); *GRHL2*, an AR coregulator (Paltoglou et al. 2017); *NR3C1*, also known as Glucocorticoid Receptor/*GR* and regulated by *AR (Xie et al. 2015)*; *PIAS1*, a competitive co-regulator of *AR (Gross et al. 2001)*; and *TLE3 (Palit et al. 2019)*, a corepressor with *AR*;
2. TFs that have been linked to castration-resistant prostate cancer: *HNF4G (Shukla et al. 2017)* and *NANOG (Jeter et al. 2011)*;
3. Putatively novel TFs without established PrCa function: *ARID1A, REPIN1*, and *VDR*.

In particular, *ARID1A, HOXB13, NR3C1* and *TLE3* were each depleted in more than 3 state categories in proximal H3K27ac asQTL peaks, but enriched in at least one distal state category, consistent with regulation specific to distal elements.

Focusing on AR, we investigated a broader landscape of AR activity measured in multiple primary prostate samples and adjacent normal tissues (Pomerantz et al. 2015) and classified into tumorspecific (TARBS) and normal-specific (NARBS) binding sites (Supplementary Figure 5). TARBS were significantly enriched in *HOXB13* and *FOXA1* asQTL peaks observed in tumors relative to balanced *HOXB13* and *FOXA1* peaks, consistent with these TFs being directly co-regulated with tumor-specific AR binding. In contrast, NARBS were significantly depleted in *HOXB13* and *FOXA1* asQTL peaks observed in normal prostate samples indicating that normal-specific AR binding is independent of these cofactors. For H3K27ac, no significant enrichment or depletion was observed for TARBS at T-asQTL peaks or NARBS at N-asQTL peaks. However, significant depletions were observed at mismatching states, i.e. NARBS were depleted at H3K27ac T-asQTL peaks, as would be expected if NARBS generally reflected lack of regulatory activity at the same site in the tumor and thus absence of asQTLs (and vice versa for TARBS). Consistent with this mechanism, no depletion of ARBS at H3K27ac asQTL peaks were observed when evaluated relative to a random baseline rather than random balanced peaks (Supplementary Figure 6).

Finally, extended our enrichment analysis to pairs of reference TFs, to investigate putative collaborative or competitive binding. For a pair of TFs *a* and *b*, we quantified the number of *a* peaks and *b* peaks that overlapped each asQTL peak, and then estimated the correlation of these two vectors as a measure of co-binding. We then performed the same quantification using random balanced peaks to estimate the difference between correlation coefficients, and significance. We identified 6 pairs of TFs with significant changes in co-binding between imbalanced and balanced peaks at an FDR of 1% (Supplementary Table 10). At *FOXA1* TvsN-asQTL peaks, the pairs *FOXA1*:*HOXB13* (change in correlation = 0.407, empirical p-value < 0.01) and *CSNK2A1*:*NANOG* (0.401, p<0.01) significantly increased in co-occurrence compared to balanced peaks. While the former are well established as co-binding factors (Pomerantz et al. 2015), the latter have not been implicated. At *HOXB13* TvsN-asQTL peaks, the pairs *H2A.Z*:*TOP1* (0.512, p < 0.01), *H2A.Z*:*E2F1* (0.354, p<0.01), *HNF4G*:*TLE3* (0.478, p<0.01), and *HNF4G*:*ARID1A* (0.462, p<0.01) all significantly increased in co-occurrence compared to balanced peaks. The histone variant *H2A.Z* has recently been implicated in activation of AR associated enhancers in prostate cancer (Valdés-Mora et al. 2017), providing a potential mechanism for increased co-binding with HOXB13 in TvsN-asQTLs. In contrast, while HNF4G has been implicated in an AR-independent transcriptional circuit in prostate cancer (Shukla et al. 2017), it has not been linked to the cobinding factors identified here. In sum, this analysis identifies extensive co-binding of TFs involved in prostate cancer regulation at tumor-specific, germline regulatory variants. Further functional validation is needed to distinguish which TF partners are necessary for the observed allelic imbalance.

## Discussion

We have demonstrated that germline allelic imbalance in somatic chromatin activity is a widespread phenomenon, and can be used to identify cis-regulatory variants from relatively modest sample sizes. asQTL peaks were significantly enriched for prostate-specific and tumorspecific eQTLs in independent datasets, serving as a validation of their regulatory function. PrCa GWAS heritability and significant associations were broadly enriched across imbalanced H3K27ac peaks relative to balanced H3K27ac peaks (a stringent baseline). 20 GWAS loci contained a fine-mapped SNP that overlapped an asQTL regulatory element and are thus putative targets for functional follow-up.

A particularly unique aspect of our study is the identification of asQTLs that were differentially active in tumors. We hypothesize that such asQTLs became newly activated in the context of widespread epigenetic reprogramming during tumorigenesis, and could be used to identify important somatic regulatory mechanisms. Indeed, we found that tumor-specific asQTL peaks were significantly enriched for multiple TFs, including both canonical PrCa TF’s like *AR* and *NKX3-1*, as well as novel PrCA-linked TF’s like *ARID1A* and *TLE3*. We additionally found H3K27ac asQTL peaks to be significantly enriched for accessible chromatin in embryonic stem cells (relative to epithelial cells), consistent with a broad process of “dedifferentiation” in tumors leading to activation of early developmental regulatory elements, whose activity is then altered by germline *cis*-regulatory variants.

Our work has several limitations. First, the ChIP-seq data we collected here primarily targeted H3K27ac and other broad histone marks, with relatively fewer individuals assayed for TF activity, thus limiting our ability to identify TF-specific mechanisms. Second, somatic copy number alterations can induce apparent allelic imbalance within carriers, and our approach accounts for this only through modelling the read count overdispersion for each sample. In the population, an imbalanced peak would only result from a somatic copy number change that is both highly recurrent and consistently aligns with a germline allele, an unlikely event. However, in our sample, peaks with a small number of germline heterozygous individuals may be biased due to somatic alterations. In the limit of a single heterozygous individual we cannot statistically distinguish between the variant or the copy number driving imbalance. Third, our analyses of asQTLs primarily relied on simple physical overlap, under the assumption that the variant driving allelic imbalance for a peak is itself likely to reside in the peak. Larger sample sizes (i.e. N>50) would provide the genetic diversity needed to perform formal statistical fine-mapping and disease colocalization in instances where this assumption does not hold. Fourth, we replicated and contextualized our analyses with publicly available epigenetic data that was typically collected in cell lines which may be systematically different from primary tissue. We have alleviated this by using a variety of datasets to avoid bias from validating in a single cell line. Our asQTL peaks have replicated with regulatory QTLs in external samples (Houlahan et al. 2019), but further application of this technique in other primary tissues would do much to demonstrate the effectiveness of this technique in identifying useful epigenetic changes.

## Conclusions

Our work demonstrates the broad utility of applying allelic imbalance methodology to detect regulatory variants that change activity across different contexts - in this case looking at differences across the normal/tumor cell state change. In particular, we have shown that such differential regulatory activity can pinpoint previously unobserved disease risk variants and implicate specific transcription factor programs. Our analyses show that we have not reached diminishing returns in terms of number of asQTLs that can be detected as a function of sample size and coverage, and we anticipate that our methodological approach will be of broad utility to studies of diverse molecular contexts for both cancer and non-cancer mechanisms.

## Methods

### Sample information

Prostate tissue was collected from 27 patients with localized primary prostate adenocarcinoma. Each patient yielded a sample of the adenocarcinoma and a sample from surrounding non-malignant prostate tissue. We performed chromatin immunoprecipitation sequencing (ChIP-seq) for H3K27ac (N=26), H3k4me2 (N=3), H3k4me3 (N=3), FOXA1 (N=5), and HOXB13 (N=5) on these samples, as well as germline SNP genotyping from blood. Germline variants were phased and imputed to the Haplotype Reference Consortium panel. Mapping and aligning was performed using *bwa;* allele-specific reads were processed according to the *WASP* pipeline (van de Geijn et al. 2015) to remove mapping bias; peaks were identified using the *MACS2* software. Allelespecific read counts were generated by the GATK ASEReadCounter (Castel et al. 2015).

### asQTL peak detection

We tested for allele-specific signal using a haplotype beta-binomial test that accounts for read overdispersion. Beta-binomial overdispersion parameters were estimated for each individual/experiment from the aligned allele-specific counts and were found to be consistently low (Normal: mean = 4.9e-4, sd = 1.35e-3; Tumour: 2.66e-3, 4.84e-3). Due to the negligible amount of overdispersion, we did not model local structural changes. For each peak and individual, haplotype-specific read counts were merged across all heterozygous read-carrying sites in the peak for a single measure of allele specificity. Every SNP within 100kb of the peak centre and carried by at least one heterozygous individual was then tested for allelic imbalance. All heterozygous individuals were tested together under the expectation of a consistent allelespecific effect. Each test was performed once for samples from normal, tumor, both, as well as a differential test between tumor and normal. Finally, peaks were considered “imbalanced” in each of these four test categories if any of the variants tested for that peak exhibited allele-specific signal at a 10% FDR. For further details of stratAS’s statistical methods and comparison to previous methods see (Gusev et al. 2019). We evaluated multiple parameter settings for asQTL detection: the size of the peaks used for testing; the distance over which we tested for SNPs around each peak and the exclusion distance between SNPs in a read. Detection sensitivity was maximized with: peaks defined as 1kb either side of the narrow peak calls derived from MACS2, and testing all SNPs within 100kb of the peak center.

### asQTL peak enrichment

To determine the general biology of this class of peaks, we measured the enrichment of these peaks for various genomic indicators of interest compared to randomly sampled balanced peaks. We applied the following workflow using bedtools (Quinlan and Hall 2010) and R:

1. For each asQTL and non-asQTL peak in that HMTF-state category, calculate how many epigenetic features of interest overlap with the peak interval, and normalise by the number of base pairs.
2. Calculate the mean number of features per base pair in the asQTL peaks (mean.imbalanced).
3. Randomly sample a number of balanced peaks equal to the number of asQTL peaks. Calculate the mean number of features per base pair in these selected non-asQTL peaks.
4. Repeat step (3) 5000 times to create a null distribution of randomly sampled balanced peaks which captures the variance in estimates of enrichment.
5. Calculate the following quantities:
  1. Mean of the balanced peaks enrichment distribution (mean.balanced)
  2. Standard deviation of balanced peaks enrichment distribution (sd)
  3. The absolute difference between mean.imbalanced and mean.balanced (balanced_to_imbalanced)
  4. The z-score of imbalanced peak enrichment: z = balanced_to_imbalanced/sd
  5. The fold enrichment: relative_enrichment = mean.imbalanced/mean.balanced
  6. The fraction rank of mean.imbalanced in the balanced peaks enrichment distribution (percentage)
  7. The empirical p (empirical_p) value:
    1. If the fraction rank is less than 0.5: empirical_p = percentage * 2
    2. If the fraction rank is more than 0.5: empirical_p = (1 - percentage) * 2

The following datasets were used in the enrichment analysis:

1. Finemapped prostate cancer GWAS SNPs. This dataset contained 3700 finemapped PrCA SNPs. To generate contiguous loci, we added a ±500kb window around each SNP and merged overlapping regions, yielding 71 loci (Dadaev et al. 2018)
2. Prostate and other tissues eQTLs from GTEx v7 (“GTEx Portal” n.d.)
3. Prostate adenocarcinoma and other cancer eQTLs from PANCAN (Gong et al. 2018)
4. ROADMAP consortium h3k27ac narrowPeak files for embryonic stem cells (Roadmap Epigenomics Consortium et al. 2015)
5. AR ChIP-seq binding sites that are differentially enriched in normal tissue (NARBS) or tumor tissue (TARBS) (Pomerantz et al. 2015)
6. H3K27ac ChIP-seq peaks in fetal prostate tissue (Pomerantz et al. 2020)

### Advanced asQTL peak enrichment

In addition to the above simple enrichment protocol, we expanded this to include a more complex protocol for the tissue-specific scores and the ES-specific score.

We complete step (1) from a basic enrichment analysis for two different features, creating two vectors of enrichment sequences *x* and *y* e.g. GTEx-prostate and GTEx-non-prostate. Our GTEx-prostate enrichment reflects the enrichment for our feature of interest, which can be split into two sources: the technical features of GTEx which affect all types of eQTL, and those which are specific to prostate eQTLs. We then calculate x/y to create one vector of the ratio between e.g. GTEx-prostate and GTEx-non-prostate enrichment. We then apply the enrichment steps (2) to (7) to calculate the standard enrichment parameters.

For the GTEx-prostate-specific score, prostate eQTLs from GTEx were used as the numerator and eQTLs from all other non-prostate tissue in GTEx was used as the denominator; for the TCGA-PRAD-specific score, PRAD eQTLs from TCGA were numerator and eQTLs from all other tumors in TCGA were denominator; for the ES-specific score, the H3K27ac peak calls for embryonic stem cells from ROADMAP were the numerator, and H3K27ac peak calls from fetal prostate cells were the denominator.

We also calculated enrichment where we change step (4) from using balance peaks as a null distribution, to using 1000 randomly shuffled intervals of equal size to the asQTL peaks as a null distribution. This is a less stringent baseline with which to compare asQTL enrichment for epigenetic features.

### Enrichment in the proximity of asQTL peaks

We quantified whether asQTL peaks exhibited unusual proximity to other asQTL peaks using random sampling. For a given pair of HMTFs *a* and *b*, we computed the mean distance between asQTL peak *a* and the nearest asQTL peak *b*. The null distribution was generated by sampling the same number of balanced peaks *a*, and computing the mean distance to the nearest asQTL peak *b*. The resampling procedure was performed multiple times to obtain the null distribution.

### Quantifying TF binding at asQTL peaks

We estimated enrichment of asQTLs at 25 transcription factors and histone marks from the LnCaP cell line from Cistrome DB (Mei et al. 2017). In addition to comparing to a background of balanced peaks, we sought to account for differences in peak coverage. For each asQTL HMTF, we constructed a background balanced peak set by matching each asQTL peak to the 20 random balanced peaks with similar read coverage. For each focal TF and asQTL peak class, we computed the number of binding sites overlapping the asQTL peaks as the numerator of the enrichment. We then randomly sampled the same number of (read matched) balanced peaks and computed the number of overlapping binding sites as the background. Enrichment was computed as the numerator over the mean of the denominator, and significance was computed relative to multiple resamplings of the denominator, using the same procedure as described for the basic enrichment analysis.

## Declarations

### Ethics approval and consent to participate

Fresh-frozen radical prostatectomy specimens were selected from the Dana–Farber Cancer Institute (DFCI) Gelb Center biobank and database as part of DFCI protocols 01-045 and 09-171 and approved by the DFCI/Harvard Cancer Center institutional review board.

### Consent for publication

Not applicable

### Availability of data and materials

All sequencing data generated for the study has been deposited in GEO (GSE130408). Full analysis outputs and summary statistics are available as Supplementary Data.

### Authors’ contributions

Each author contributed to conception (AS, MLF, AG); design of the work (AS, MLF, AG); acquisition, analysis, and interpretation of data (all authors); and drafting the manuscript (AS, MLF, AG).

## Acknowledgements

The authors would like to acknowledge, Claudia Giambartolomei, Cynthia Kalita, Bogdan Pasaniuc, Austin Wang, Douglas Yao, and the members of the Freedman and Gusev labs for helpful discussions.

## Supplementary Figures

Supplementary Figure 1: PCA of all peaks and asQTL peaks only

Supplementary Figure 2: Distribution of TvsN-asQTL peaks depending on the allelic fraction of their associated SNPs in normal and tumour tissue samples

Supplementary Figure 3: Enrichment of HMTF-state asQTL peaks for 6 genetic and epigenetic features with a background of random genomic intervals instead of balanced peaks

Supplementary Figure 4: Enrichment of HMTF-state asQTL peaks for ES-specific enrichment with a category of TvsN-asQTL and T-asQTL peaks combined

Supplementary Figure 5: Enrichment of HMTF-state peaks for AR binding sites in tumour cells, normal prostate cells and in the LnCaP cell

Supplementary Figure 6: Enrichment of HMTF-state peaks for AR binding sites in tumour cells, normal prostate cells and in the LnCaP cell against a background of random genomic intervals instead of balanced peaks

## Tables

Table 1: Number of samples and peaks by HMTF-state combination

Table 2: Number of GWAS loci lying within different HMTF-state categories of asQTL peaks

## Supplementary Tables

Supplementary Table 1: The individual samples used for the analysis, and what HMTF was used (the same sample may be assessed for multiple HMTFs)

Supplementary Table 2: The correlation in allelic fraction for asQTL SNPs between different HMTFs

Supplementary Table 3: The median absolute distance between asQTL peaks of one state/HMTF combination and the asQTL/any peak of another HMTF-state combination using standard enrichment procedure

Supplementary Table 4: LDSC regression results for imbalanced and balanced peak HMTF-state categories

Supplementary Table 5: Full results of enrichment of HMTF-state peaks for 6 genetic and epigenetic features with a background of random genomic intervals instead of balanced peaks

Supplementary Table 6: List of 16 fine-mapped SNPs who lie within the overlap intervals between PrCA GWAS loci and imbalanced *HOXB13* and *FOXA1* peaks, and not imbalanced h3k27ac, h3k4me3, h3k4me2

Supplementary Table 7: Full results of enrichment of HMTF-state peaks for 6 genetic and epigenetic features with a background of balanced peaks, providing a conservative estimate of enrichment

Supplementary Table 8: Enrichments of 25 epigenetic features from ROADMAP within our asQTL peaks with a conservative balanced peaks background

Supplementary Table 9: Enrichments of 25 epigenetic features from ROADMAP within H3K27ac peaks stratified by distance to the TSS

Supplementary Table 10: Change in correlation between pairs TFs co-located within asQTL peaks against randomly sampled balanced peaks, matched for read coverage

## References

Barbeira, Alvaro N., Scott P. Dickinson, Rodrigo Bonazzola, Jiamao Zheng, Heather E. Wheeler, Jason M. Torres, Eric S. Torstenson, et al. 2018. “Exploring the Phenotypic Consequences of Tissue Specific Gene Expression Variation Inferred from GWAS Summary Statistics.”Nature Communications 9 (1): 1825.

Benaglio, Paola, Agnieszka D’Antonio-Chronowska, Wubin Ma, Feng Yang, William W. Young Greenwald, Margaret K. R. Donovan, Christopher DeBoever, et al. 2019. “Allele-Specific NKX2-5 Binding Underlies Multiple Genetic Associations with Human Electrocardiographic Traits.”Nature Genetics 51 (10): 1506–17.

Bhatia-Gaur, R., A. A. Donjacour, P. J. Sciavolino, M. Kim, N. Desai, P. Young, C. R. Norton, et al. 1999. “Roles for Nkx3.1 in Prostate Development and Cancer.”Genes & Development 13 (8): 966–77.

Brawn, P. N. 1983. “The Dedifferentiation of Prostate Carcinoma.”Cancer 52 (2): 246–51.

Castel, Stephane E., Ami Levy-Moonshine, Pejman Mohammadi, Eric Banks, and Tuuli Lappalainen. 2015. “Tools and Best Practices for Data Processing in Allelic Expression Analysis.”Genome Biology 16 (September): 195.

Conti, David V., Burcu F. Darst, Lilit C. Moss, Edward J. Saunders, Xin Sheng, Alisha Chou, Fredrick R. Schumacher, et al. 2021. “Trans-Ancestry Genome-Wide Association Meta-Analysis of Prostate Cancer Identifies New Susceptibility Loci and Informs Genetic Risk Prediction.”Nature Genetics 53 (1): 65–75.

Dadaev, Tokhir, Edward J. Saunders, Paul J. Newcombe, Ezequiel Anokian, Daniel A. Leongamornlert, Mark N. Brook, Clara Cieza-Borrella, et al. 2018. “Fine-Mapping of Prostate Cancer Susceptibility Loci in a Large Meta-Analysis Identifies Candidate Causal Variants.”Nature Communications 9 (1): 2256.

Emami, Nima C., Linda Kachuri, Travis J. Meyers, Rajdeep Das, Joshua D. Hoffman, Thomas J. Hoffmann, Donglei Hu, et al. 2019. “Association of Imputed Prostate Cancer Transcriptome with Disease Risk Reveals Novel Mechanisms.”Nature Communications 10 (1): 3107.

Gamazon, Eric R., GTEx Consortium, Heather E. Wheeler, Kaanan P. Shah, Sahar V. Mozaffari, Keston Aquino-Michaels, Robert J. Carroll, et al. 2015. “A Gene-Based Association Method for Mapping Traits Using Reference Transcriptome Data.”Nature Genetics. https://doi.org/10.1038/ng.3367.

Gate, Rachel E., Christine S. Cheng, Aviva P. Aiden, Atsede Siba, Marcin Tabaka, Dmytro Lituiev, Ido Machol, et al. 2018. “Genetic Determinants of Co-Accessible Chromatin Regions in Activated T Cells across Humans.”Nature Genetics 50 (8): 1140–50.

Geeleher, Paul, Aritro Nath, Fan Wang, Zhenyu Zhang, Alvaro N. Barbeira, Jessica Fessler, Robert L. Grossman, Cathal Seoighe, and R. Stephanie Huang. 2018. “Cancer Expression Quantitative Trait Loci (eQTLs) Can Be Determined from Heterogeneous Tumor Gene Expression Data by Modeling Variation in Tumor Purity.”Genome Biology. https://doi.org/10.1186/s13059-018-1507-0.

Geijn, Bryce van de, Graham McVicker, Yoav Gilad, and Jonathan K. Pritchard. 2015. “WASP: Allele-Specific Software for Robust Molecular Quantitative Trait Locus Discovery.”Nature Methods 12 (11): 1061–63.

Gong, Jing, Shufang Mei, Chunjie Liu, Yu Xiang, Youqiong Ye, Zhao Zhang, Jing Feng, et al. 2018. “PancanQTL: Systematic Identification of Cis-eQTLs and Trans-eQTLs in 33 Cancer Types.”Nucleic Acids Research 46 (D1): D971–76.

Gross, M., B. Liu, J. Tan, F. S. French, M. Carey, and K. Shuai. 2001. “Distinct Effects of PIAS Proteins on Androgen-Mediated Gene Activation in Prostate Cancer Cells.”Oncogene 20 (29): 3880–87.

Grubert, Fabian, Judith B. Zaugg, Maya Kasowski, Oana Ursu, Damek V. Spacek, Alicia R. Martin, Peyton Greenside, et al. 2015. “Genetic Control of Chromatin States in Humans Involves Local and Distal Chromosomal Interactions.”Cell 162 (5): 1051–65.

“GTEx Portal.” n.d. Accessed April 19, 2021. https://www.gtexportal.org/home/faq.

Gusev, Alexander, Arthur Ko, Huwenbo Shi, Gaurav Bhatia, Wonil Chung, Brenda W. J. H. Penninx, Rick Jansen, et al. 2016. “Integrative Approaches for Large-Scale Transcriptome-Wide Association Studies.”Nature Genetics 48 (3): 245–52.

Gusev, Alexander, Huwenbo Shi, Gleb Kichaev, Mark Pomerantz, Fugen Li, Henry W. Long, Sue A. Ingles, et al. 2016. “Atlas of Prostate Cancer Heritability in European and African-American Men Pinpoints Tissue-Specific Regulation.”Nature Communications 7 (April): 10979.

Gusev, Alexander, Sandor Spisak, Andre P. Fay, Hallie Carol, Kevin C. Vavra, Sabina Signoretti, Viktoria Tisza, et al. 2019. “Allelic Imbalance Reveals Widespread Germline-Somatic Regulatory Differences and Prioritizes Risk Loci in Renal Cell Carcinoma.”bioRxiv. https://doi.org/10.1101/631150.

Houlahan, Kathleen E., Yu-Jia Shiah, Alexander Gusev, Jiapei Yuan, Musaddeque Ahmed, Anamay Shetty, Susmita G. Ramanand, et al. 2019. “Genome-Wide Germline Correlates of the Epigenetic Landscape of Prostate Cancer.”Nature Medicine 25 (10): 1615–26.

Jeter, C. R., B. Liu, X. Liu, X. Chen, C. Liu, T. Calhoun-Davis, J. Repass, H. Zaehres, J. J. Shen, and D. G. Tang. 2011. “NANOG Promotes Cancer Stem Cell Characteristics and Prostate Cancer Resistance to Androgen Deprivation.”Oncogene 30 (36): 3833–45.

Knowles, David A., Joe R. Davis, Hilary Edgington, Anil Raj, Marie-Julie Favé, Xiaowei Zhu, James B. Potash, et al. 2017. “Allele-Specific Expression Reveals Interactions between Genetic Variation and Environment.”Nature Methods 14 (7): 699–702.

Kumasaka, Natsuhiko, Andrew J. Knights, and Daniel J. Gaffney. 2016. “Fine-Mapping Cellular QTLs with RASQUAL and ATAC-Seq.”Nature Genetics 48 (2): 206–13.

Kumasaka, Natsuhiko, Andrew J. Knights, and Daniel J. Gaffney. 2019. “High-Resolution Genetic Mapping of Putative Causal Interactions between Regions of Open Chromatin.”Nature Genetics 51 (1): 128–37.

Link, Verena M., Sascha H. Duttke, Hyun B. Chun, Inge R. Holtman, Emma Westin, Marten A. Hoeksema, Yohei Abe, et al. 2018. “Analysis of Genetically Diverse Macrophages Reveals Local and Domain-Wide Mechanisms That Control Transcription Factor Binding and Function.”Cell 173 (7): 1796–1809.e17.

Liu, Tao, Jorge A. Ortiz, Len Taing, Clifford A. Meyer, Bernett Lee, Yong Zhang, Hyunjin Shin, et al. 2011. “Cistrome: An Integrative Platform for Transcriptional Regulation Studies.”Genome Biology 12 (8): R83.

Liu, Xuanyao, Yang I. Li, and Jonathan K. Pritchard. 2019. “Trans Effects on Gene Expression Can Drive Omnigenic Inheritance.”Cell 177 (4): 1022–34.e6.

Mancuso, Nicholas, Simon Gayther, Alexander Gusev, Wei Zheng, Kathryn L. Penney, Zsofia Kote-Jarai, Rosalind Eeles, et al. 2018. “Large-Scale Transcriptome-Wide Association Study Identifies New Prostate Cancer Risk Regions.”Nature Communications 9 (1): 4079.

Maurano, Matthew T., Eric Haugen, Richard Sandstrom, Jeff Vierstra, Anthony Shafer, Rajinder Kaul, and John A. Stamatoyannopoulos. 2015. “Large-Scale Identification of Sequence Variants Influencing Human Transcription Factor Occupancy in Vivo.”Nature Genetics 47 (12): 1393–1401.

Mei, Shenglin, Qian Qin, Qiu Wu, Hanfei Sun, Rongbin Zheng, Chongzhi Zang, Muyuan Zhu, et al. 2017. “Cistrome Data Browser: A Data Portal for ChIP-Seq and Chromatin Accessibility Data in Human and Mouse.”Nucleic Acids Research 45 (D1): D658–62.

Palit, Sander Al, Daniel Vis, Suzan Stelloo, Cor Lieftink, Stefan Prekovic, Elise Bekers, Ingrid Hofland, et al. 2019. “TLE3 Loss Confers AR Inhibitor Resistance by Facilitating GR-Mediated Human Prostate Cancer Cell Growth.”eLife 8 (December). https://doi.org/10.7554/eLife.47430.

Paltoglou, Steve, Rajdeep Das, Scott L. Townley, Theresa E. Hickey, Gerard A. Tarulli, Isabel Coutinho, Rayzel Fernandes, et al. 2017. “Novel Androgen Receptor Coregulator GRHL2 Exerts Both Oncogenic and Antimetastatic Functions in Prostate Cancer.”Cancer Research 77 (13): 3417–30.

Polak, Paz, Rosa Karlić, Amnon Koren, Robert Thurman, Richard Sandstrom, Michael Lawrence, Alex Reynolds, et al. 2015. “Cell-of-Origin Chromatin Organization Shapes the Mutational Landscape of Cancer.”Nature 518 (7539): 360–64.

Pomerantz, Mark M., Fugen Li, David Y. Takeda, Romina Lenci, Apurva Chonkar, Matthew Chabot, Paloma Cejas, et al. 2015. “The Androgen Receptor Cistrome Is Extensively Reprogrammed in Human Prostate Tumorigenesis.”Nature Genetics 47 (11): 1346–51.

Pomerantz, Mark M., Xintao Qiu, Yanyun Zhu, David Y. Takeda, Wenting Pan, Sylvan C. Baca, Alexander Gusev, et al. 2020. “Prostate Cancer Reactivates Developmental Epigenomic Programs during Metastatic Progression.”Nature Genetics 52 (8): 790–99.

Quinlan, Aaron R., and Ira M. Hall. 2010. “BEDTools: A Flexible Suite of Utilities for Comparing Genomic Features.”Bioinformatics 26 (6): 841–42.

Reshef, Yakir A., Hilary K. Finucane, David R. Kelley, Alexander Gusev, Dylan Kotliar, Jacob C. Ulirsch, Farhad Hormozdiari, et al. 2018. “Detecting Genome-Wide Directional Effects of Transcription Factor Binding on Polygenic Disease Risk.”Nature Genetics 50 (10): 1483–93.

Roadmap Epigenomics Consortium, Anshul Kundaje, Wouter Meuleman, Jason Ernst, Misha Bilenky, Angela Yen, Alireza Heravi-Moussavi, et al. 2015. “Integrative Analysis of 111 Reference Human Epigenomes.”Nature 518 (7539): 317–30.

Schumacher, Fredrick R., Ali Amin Al Olama, Sonja I. Berndt, Sara Benlloch, Mahbubl Ahmed, Edward J. Saunders, Tokhir Dadaev, et al. 2018. “Association Analyses of More than 140,000 Men Identify 63 New Prostate Cancer Susceptibility Loci.”Nature Genetics 50 (7): 928–36.

Shukla, Shipra, Joanna Cyrta, Devan A. Murphy, Edward G. Walczak, Leili Ran, Praveen Agrawal, Yuanyuan Xie, et al. 2017. “Aberrant Activation of a Gastrointestinal Transcriptional Circuit in Prostate Cancer Mediates Castration Resistance.”Cancer Cell 32 (6): 792–806.e7.

Sunkel, Benjamin, Dayong Wu, Zhong Chen, Chiou-Miin Wang, Xiangtao Liu, Zhenqing Ye, Aaron M. Horning, et al. 2016. “Integrative Analysis Identifies Targetable CREB1/FoxA1 Transcriptional Co-Regulation as a Predictor of Prostate Cancer Recurrence.”Nucleic Acids Research 44 (9): 4105–22.

Tomlins, Scott A., Daniel R. Rhodes, Sven Perner, Saravana M. Dhanasekaran, Rohit Mehra, Xiao-Wei Sun, Sooryanarayana Varambally, et al. 2005. “Recurrent Fusion of TMPRSS2 and ETS Transcription Factor Genes in Prostate Cancer.”Science 310 (5748): 644–48.

Valdés-Mora, Fátima, Cathryn M. Gould, Yolanda Colino-Sanguino, Wenjia Qu, Jenny Z. Song, Kylie M. Taylor, Fabian A. Buske, et al. 2017. “Acetylated Histone Variant H2A.Z Is Involved in the Activation of Neo-Enhancers in Prostate Cancer.”Nature Communications 8 (1): 1346.

Waszak, Sebastian M., Olivier Delaneau, Andreas R. Gschwind, Helena Kilpinen, Sunil K. Raghav, Robert M. Witwicki, Andrea Orioli, et al. 2015. “Population Variation and Genetic Control of Modular Chromatin Architecture in Humans.”Cell 162 (5): 1039–50.

Whitington, Thomas, Ping Gao, Wei Song, Helen Ross-Adams, Alastair D. Lamb, Yuehong Yang, Ilaria Svezia, et al. 2016. “Gene Regulatory Mechanisms Underpinning Prostate Cancer Susceptibility.”Nature Genetics 48 (4): 387–97.

Xie, Ning, Helen Cheng, Dong Lin, Liangliang Liu, Ou Yang, Li Jia, Ladan Fazli, et al. 2015. “The Expression of Glucocorticoid Receptor Is Negatively Regulated by Active Androgen Receptor Signaling in Prostate Tumors.”International Journal of Cancer. Journal International Du Cancer 136 (4): E27–38.

