## Supplementary figures and images for "Allele-specific epigenetic activity in prostate cancer and normal prostate tissue implicates prostate cancer risk mechanisms"

### Supplemental Figure 1

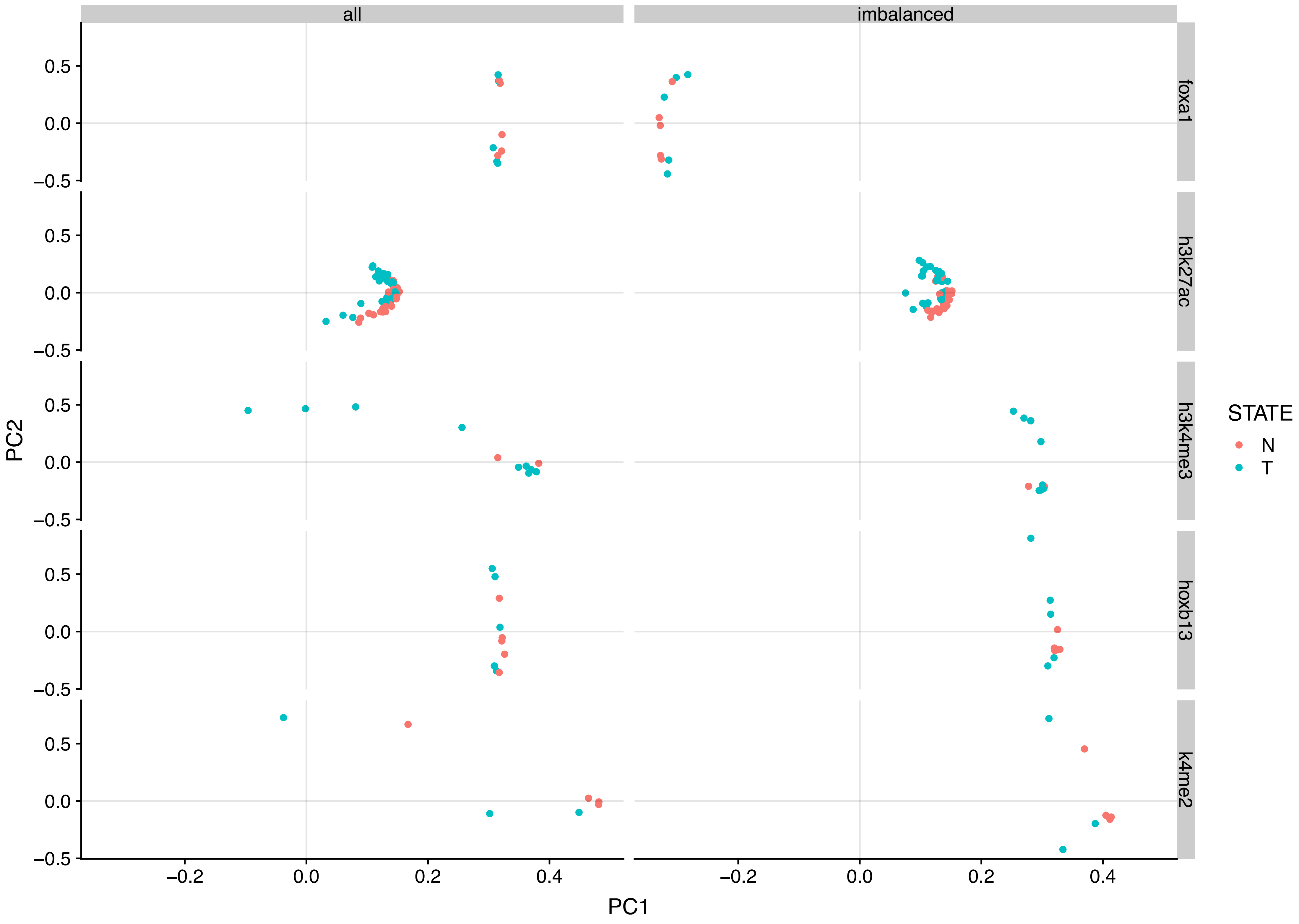

### Supplemental Figure 2

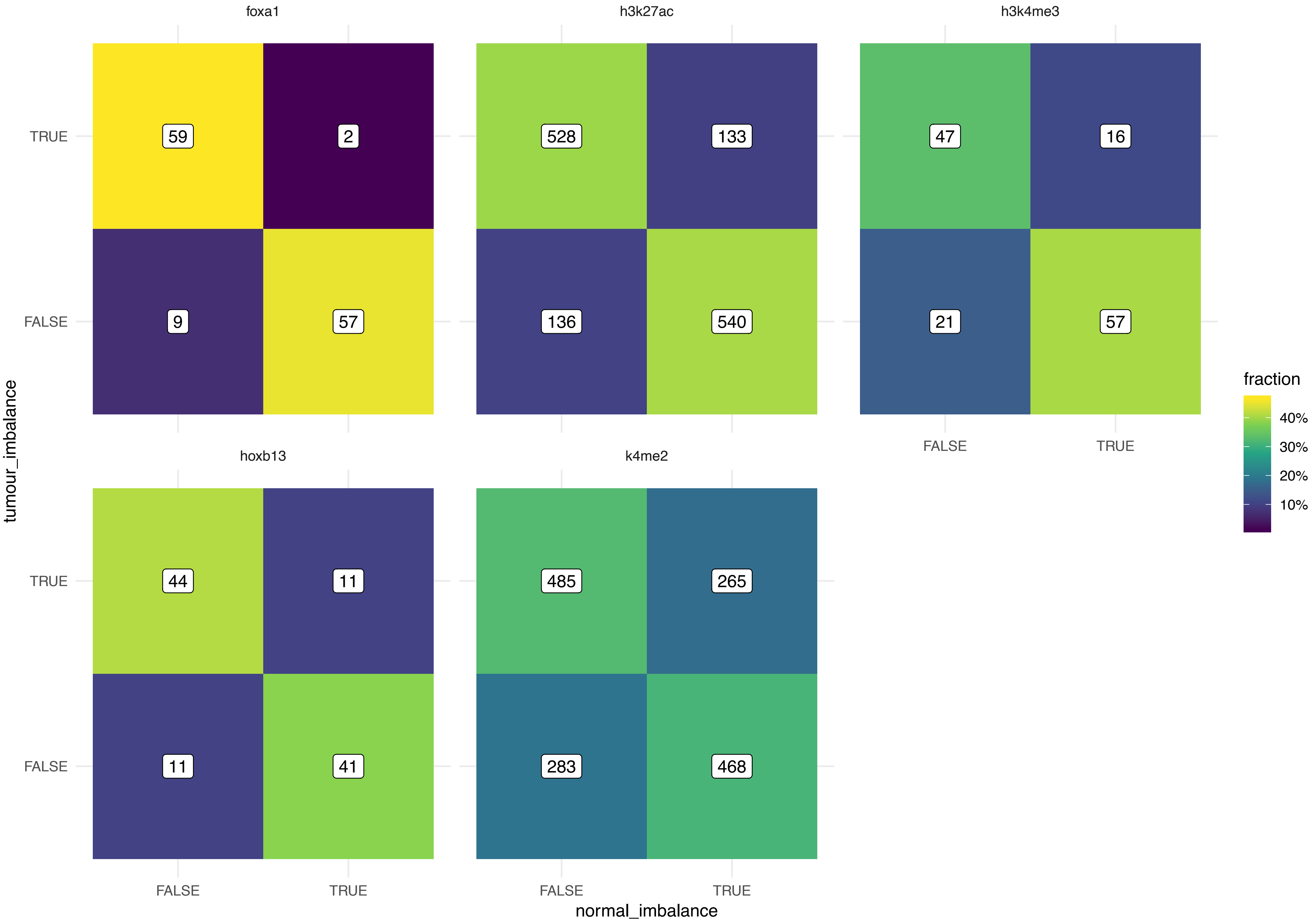

### Supplemental Figure 3

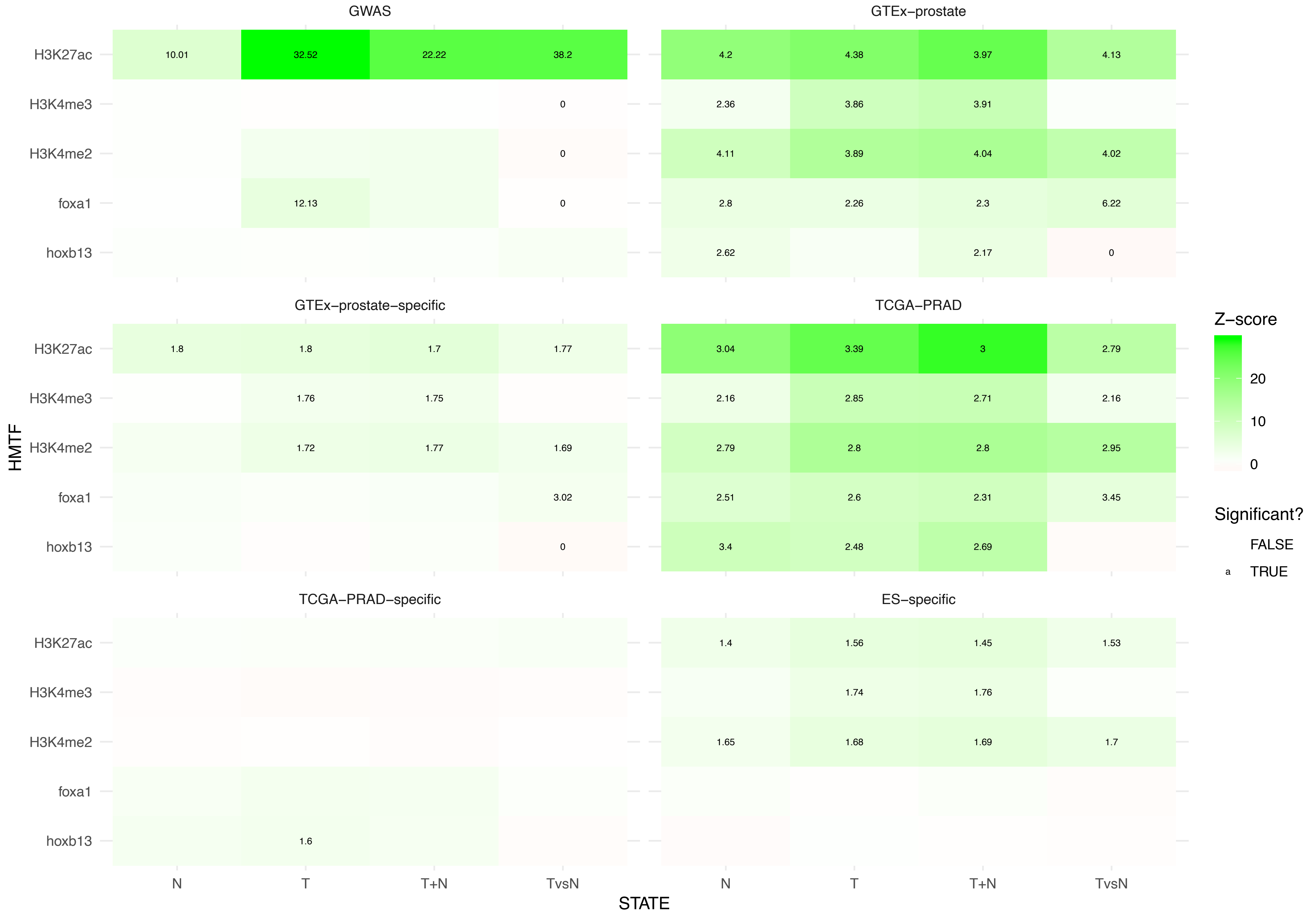

### Supplemental Figure 4

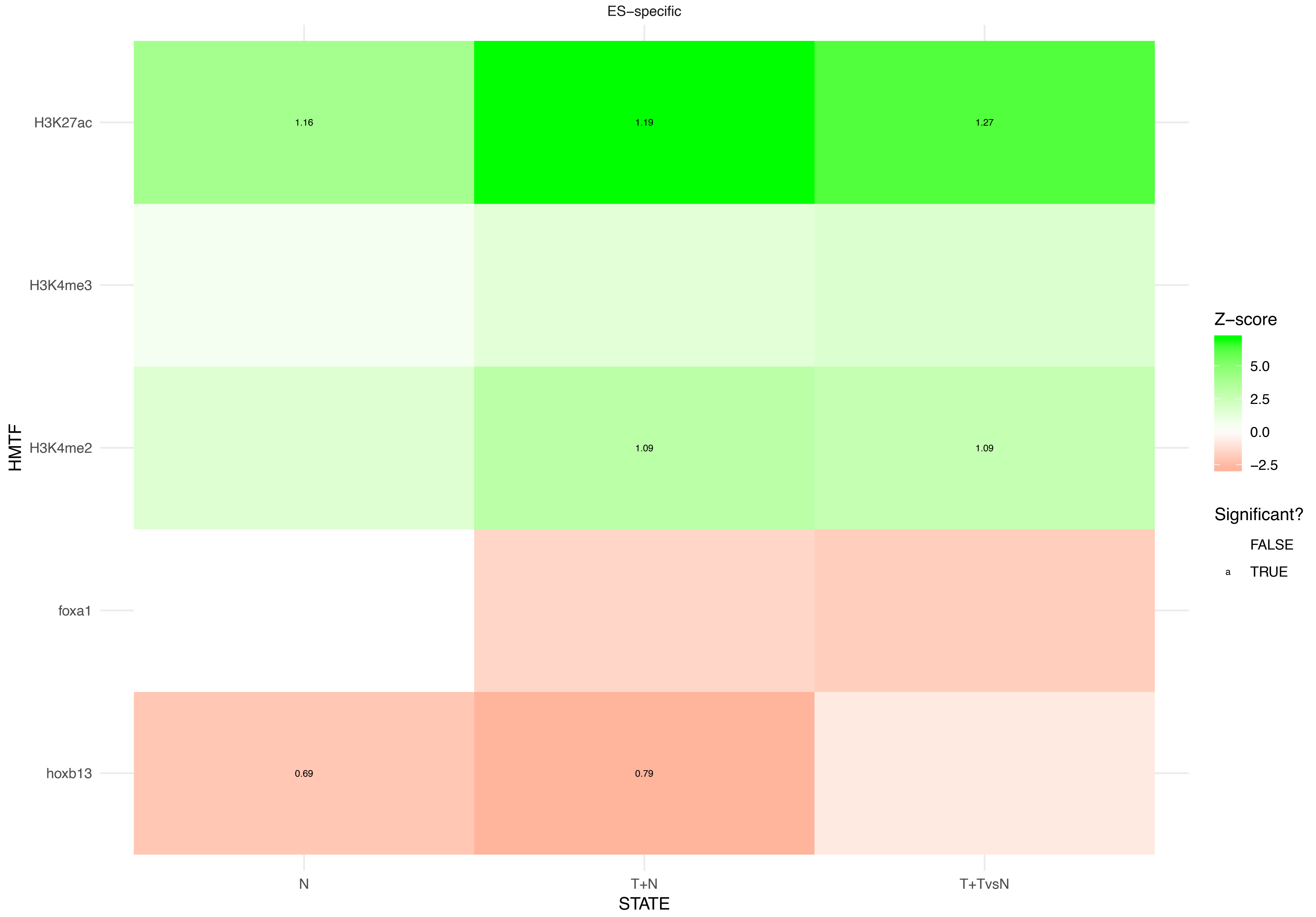

### Supplemental Figure 5

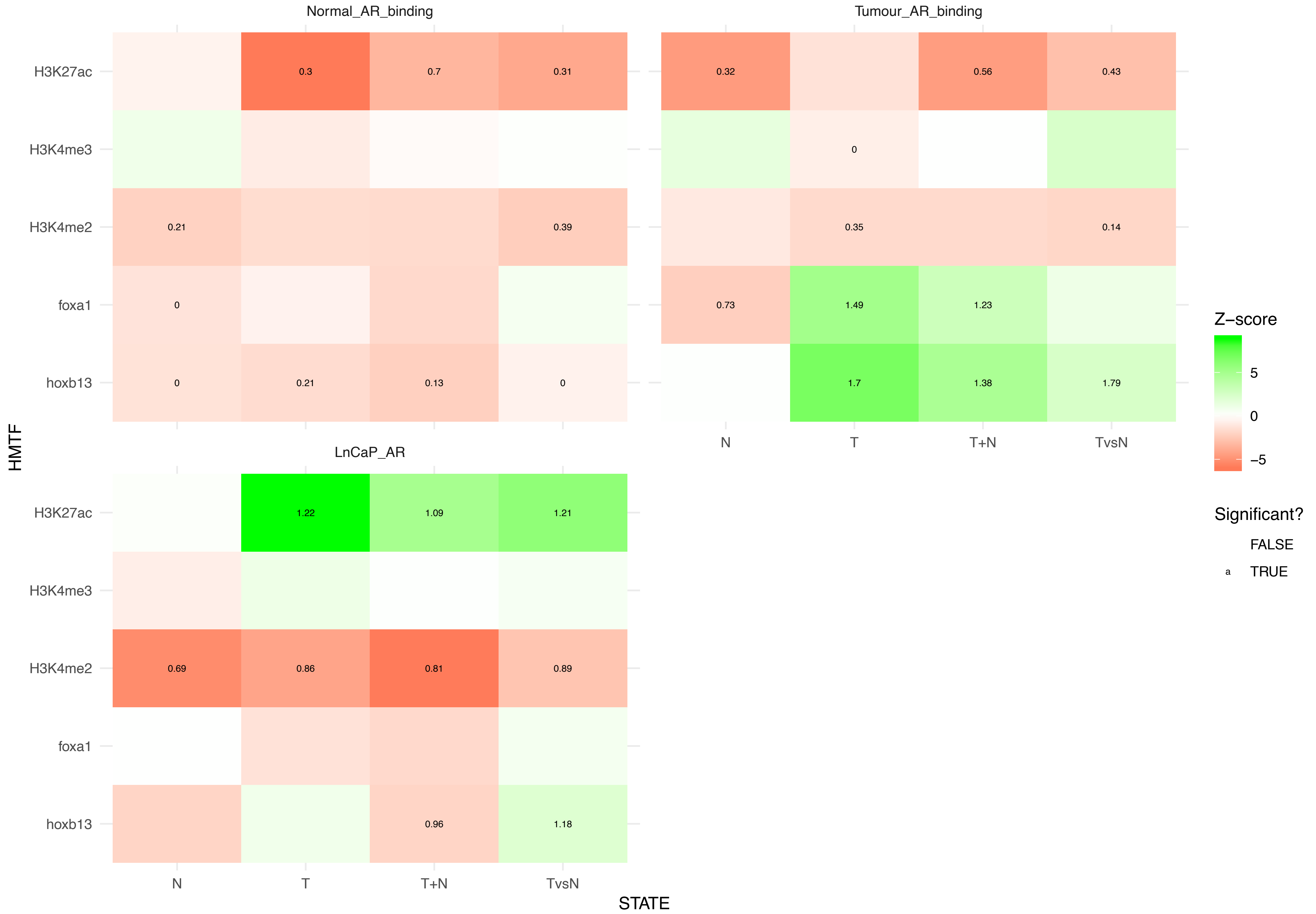

### Supplemental Figure 6

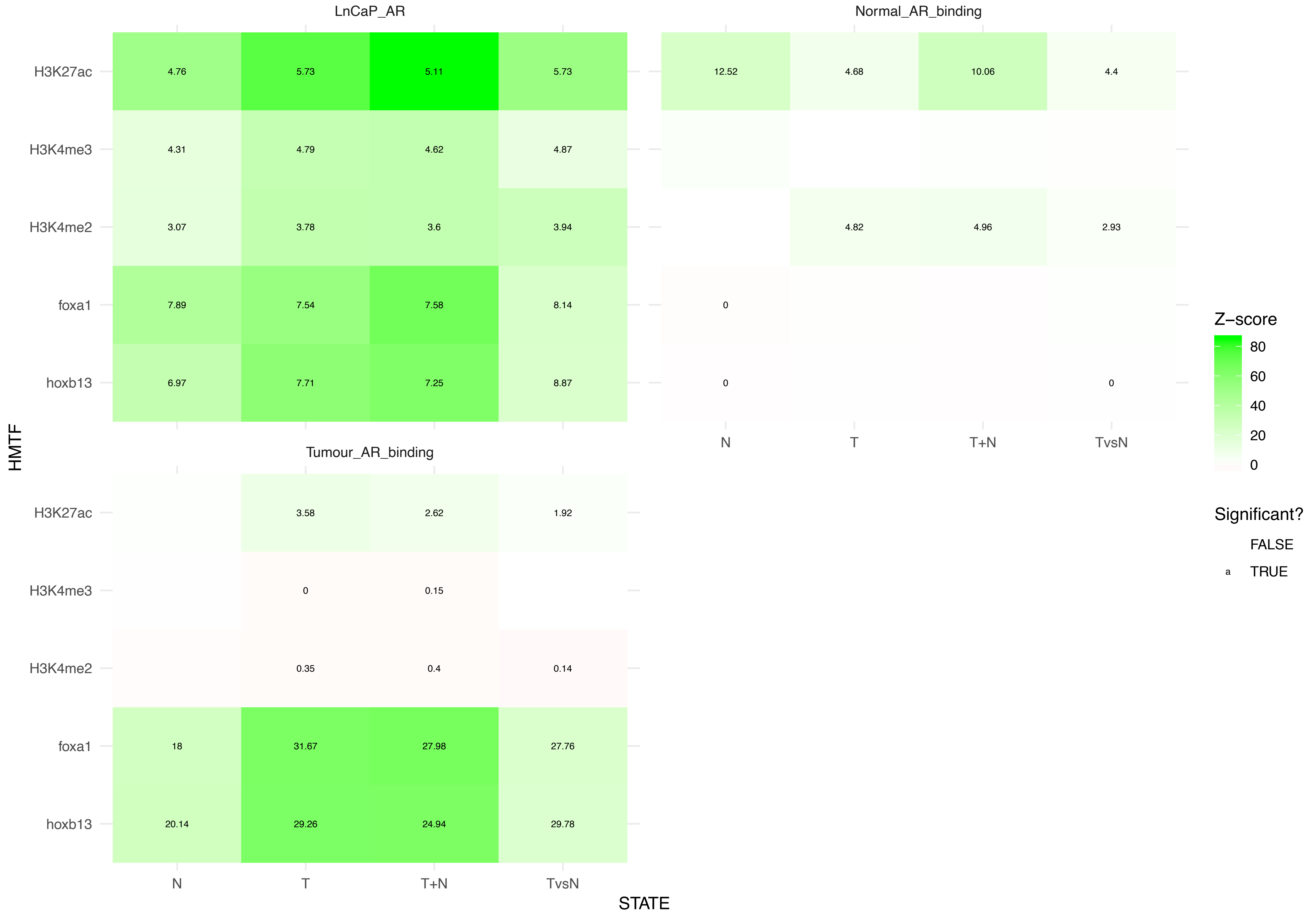
